# Hematopoietic Stem Cell Dynamics are Regulated by Progenitor Demand: Lessons from a Quantitative Modeling Approach

**DOI:** 10.1101/305367

**Authors:** Markus Klose, Maria Carolina Florian, Hartmut Geiger, Ingmar Glauche

## Abstract

The prevailing view on murine hematopoiesis and on hematopoietic stem cells (HSC) in particular derives from experiments that are related to regeneration after irradiation and HSC transplantation. However, over the past years, different experimental techniques have been developed to investigate hematopoiesis under homeostatic conditions, thereby providing access to proliferation and differentiation rates of hematopoietic stem and progenitor cells in the unperturbed situation. Moreover, it has become clear that hematopoiesis undergoes distinct changes during aging with large effects on HSC abundance, lineage contribution, asymmetry of division and self-renewal potential. However, it is currently not fully resolved how stem and progenitor cells interact to respond to varying demands and how this balance is altered by an aging-induced shift in HSC polarity.

Here, we present an *in-silico* model to investigate the dynamics of HSC response to varying demand. By introducing an internal feedback between stem and progenitor cells, the model is suited to consistently describe both hematopoietic homeostasis and regeneration, including the limited regulation of HSCs in the homeostatic situation. The model further explains the age-dependent increase in phenotypic HSCs as a consequence of the cells’ inability to preserve divisional asymmetry.

Our model suggests a dynamically regulated population of intrinsically asymmetrically dividing HSCs as suitable control mechanism that adheres with many qualitative and quantitative findings on hematopoietic recovery after stress and aging. The modeling approach thereby illustrates how a mathematical formalism can support the conceptual and the quantitative understanding of regulatory principles in HSC biology.

## Introduction

Regenerating tissues contain stem cells as source for tissue replacement and repair. In the blood system, long-term hematopoietic stem cells (LT-HSCs) have been characterized as a rare cell population with usually slow turnover. The turnover can be dramatically accelerated in stress situations such as cytotoxic treatments or post transplantation (Moore and Lemischka, 2006; Morrison and Weissman, 1994; Wichmann and Loeffler, 1985; Wilson et al., 2008). Most of the findings on the function of LT-HSCs have been derived in conditions in which HSCs were transplanted. In fact, regeneration of the hematopoietic system after cell transplantation into an irradiated host is still considered to be the gold standard for a functional definition of LT-HSCs. Minimally invasive methods for marking of LT-HSCs (Busch et al., 2015; Rodriguez-Fraticelli et al., 2018; Schoedel et al., 2016; Sun et al., 2014; Yu et al., 2016) recently allowed studying the hematopoietic system in unperturbed, i.e. not transplanted, settings, thereby adding new insights into the functional role of LT-HSCs and their progenitor populations. Based on these new findings, it appears that multipotent progenitors ensure the long-term supply with functional, differentiated hematopoietic cells in the homeostatic situation, while rarely depending on the turnover of the LT-HSC population. The question remains how such a stable regenerating system reacts to an increased cell demand, for example in case of severe perturbations.

It has also been shown that the hematopoietic system undergoes dramatic changes with age, most prominently characterized by an increasing contribution towards myeloid cell types at the expense of lymphoid differentiation, the overall loss of regenerative potential and an increase in the number of phenotypically defined LT-HSCs (Beerman et al., 2010; Chambers et al., 2007; de Haan and Van Zant, 1999; Dykstra et al., 2011; Florian et al., 2012; Geiger et al., 2013; Henry et al., 2011; Morrison et al., 1996; Noda et al., 2009; Rossi et al., 2005; Sudo et al., 2000). A single, causative reason for this aging phenotype has not been identified yet, although different mechanisms have been postulated such as accumulating DNA damage, increasing ROS levels or telomere shortening (Armanios et al., 2009; Geiger et al., 2013; Geiger et al., 2014; Henry et al., 2011; Rossi et al., 2007; Rossi et al., 2005). Recently, it became increasingly prominent that especially the declining integrity of epigenetic signatures in stem and progenitor cells might be directly involved in the functional deregulation of aged LT-HSCs. The potential reasons for this epigenetic aging are manifold. Previous findings further identified a tight link between the regulation of cell polarity and distinct epigenetic marks (Florian et al., 2012). Most importantly, it could be shown that polarity was lost with age, thereby leading to more apolar cells that will undergo symmetric cell divisions (Florian et al., 2018). The consequences of such a shift in the ability to undergo asymmetric cell divisions on the level of the population of LT-HSCs is unknown.

Hematopoietic regeneration is usually depicted as a hierarchical process in which LT-HSCs continuously contribute to a downstream, subsequently amplifying set of cell compartments. Experimental results on the role of LT-HSC turnover in both the homeostatic and the challenged situation are essential to estimate the contribution of individual cell populations to the overall hematopoietic maintenance under changing demands as well as upon aging. Mathematical formalisms are essential to derive interpretable approximations of quantities that are not experimentally accessible yet, such as rates of HSC turnover (Busch et al., 2015; Glauche et al., 2009; Wilson et al., 2008) or the “flux” between cell populations. Moreover, beyond supporting the data analysis, mathematical models represent essential tools to translate biological hypotheses into quantitative and testable predictions. Such models are instrumental to speculate about functional mechanisms and to provide an unbiased and minimalistic view on the role of regulatory principles in hematopoiesis.

We developed a mathematical model of hematopoietic stem cell organization that explicitly accounts for the ambivalence between hematopoietic maintenance in the unchallenged homeostatic situation and the immediate stress response, while intrinsically representing LT-HSC divisions as asymmetric events. Our model establishes a unifying framework which reproduces both the rare contribution and the slow recovery of HSCs in the homeostatic situation (Busch et al., 2015; Schoedel et al., 2016; Sheikh et al., 2016; Sun et al., 2014), as well as the accelerated responses after perturbation (see references in (Wichmann and Loeffler, 1985)). The model extends to address the apparent increase in phenotypic LT-HSCs as a consequence of an impaired HSC control due to the loss of cell polarity and the consequent inability of the cells to divide asymmetrically (Florian et al., 2018). Our model suggests a dynamically regulated population of intrinsically asymmetrically dividing LT-HSCs as suitable control mechanism that adheres with many qualitative and quantitative findings on hematopoietic recovery after stress and aging.

## Methods

### Model setup

We present a simple mathematical model based on ordinary differential equations (ODEs) describing proliferation and differentiation of LT-HSC, short-term hematopoietic stem cell (ST-HSC) and multipotent progenitor (MPP) populations (Fig. 1a). While the qualitative features of hematopoietic stem cell regulation can be represented by a LT-HSC and ST-HSC compartment only, we extend our model to better describe available experimental data by adding a third (MPP) compartment. The model has the following features:

- **Turnover and repopulation.** All cells are able to proliferate with respective rates *p* and differentiate into downstream compartments. We do not account for apoptotic events. Additionally, the effective proliferation of LT-HSCs *p*_LT, eff_ is regulated according to the *demand* within the ST-HSC compartment. Technically, this is implemented as a logistic growth limitation, where 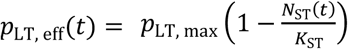 with a carrying capacity *K*_*ST*_.
- **Symmetry of LT-HSC division.** For LT-HSCs, we consider symmetric divisions (with rate *s*, ranging between 0 and 1), in which both daughter cells retain the LT-HSC identity, and asymmetric divisions (with rate 1 – *s*), in which one daughter remains a stem cell, whereas the other daughter cell instantly differentiates into the downstream ST-HSC compartment. We neglect the possibility of symmetric divisions with both daughters immediately differentiating. To account for division-independent differentiation of LT-HSCs, we introduce a parameter *d*_0_ describing a simple additional flux from the LT-HSC to the ST-HSC compartment.
- **Aging.** We account for an age-dependent change of the fraction of symmetric cell divisions by explicitly formulating the symmetry parameter *s* = *s*(*t*) as a time-dependent quantity.

The resulting ODE describes the dynamics of the LT-HSC compartment in terms of cell numbers *N*_*LT*_(*t*):

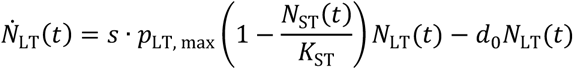

**Figure 1:**
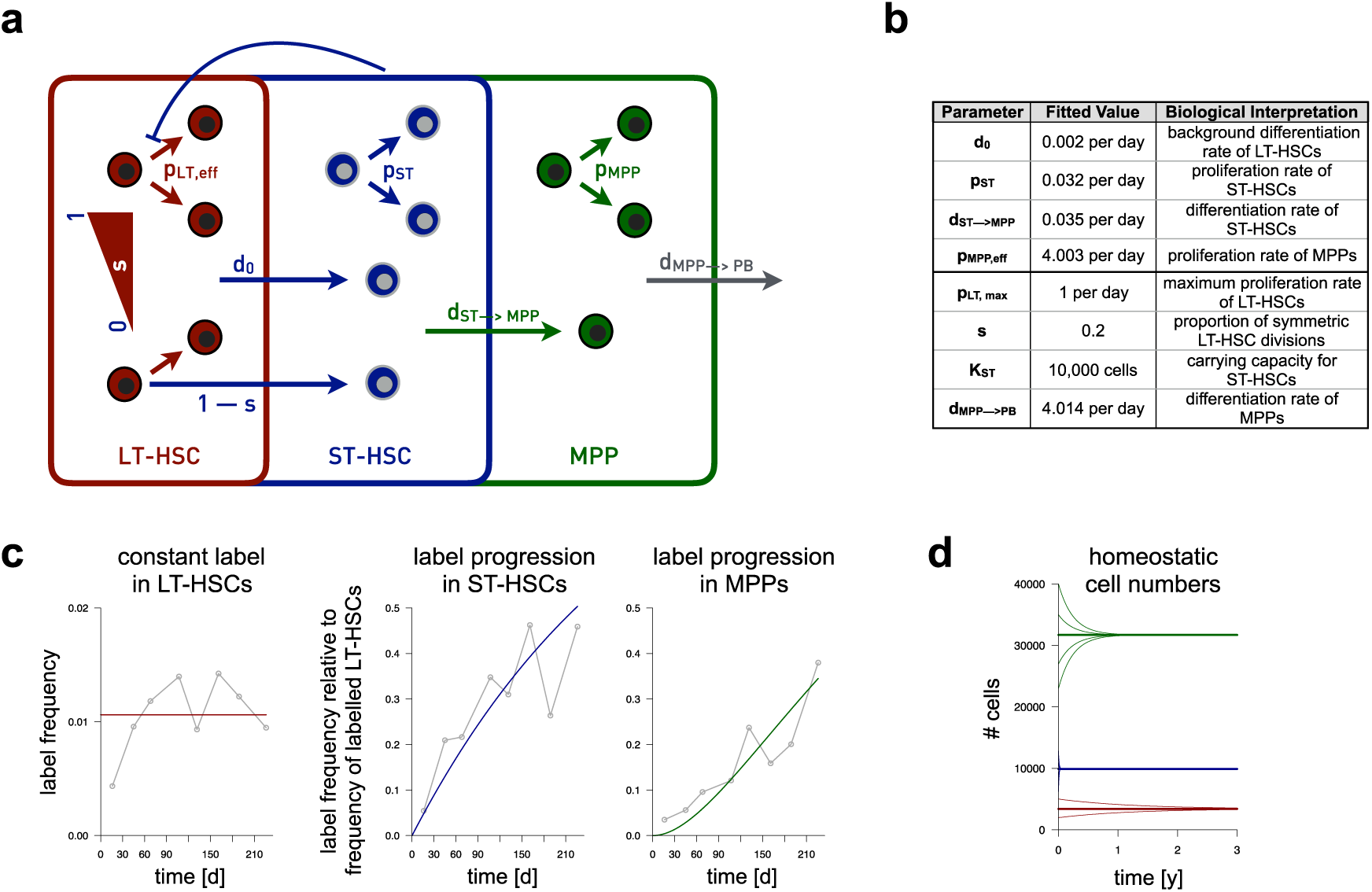
Model setup and parameter estimation. **(a)** Sketch of the mathematical model setup. LT-HSCs (red) are able to proliferate with an effective proliferation rate *p*_LT, eff_ which is regulated by ST-HSC demand. LT-HSCs differentiate instantly after division according to symmetry parameter 0 < *s* < 1. Additionally, the model incorporates a background differentiation of *d*_0_. ST-HSCs (blue) and MPPs (green) are able to proliferate and differentiate with respective rates p and *d*. Additionally, there is an influx from the respective upstream compartment. **(b)** Table of fitted values for estimated (upper part) and assigned (lower part) parameter values. **(c)** Label progression time courses for the homeostatic parameter set in **(b)** for LT-HSCs (red), ST-HSCs (blue) and MPPs (green), respectively. Grey curves correspond to label progression data by (Busch et al., 2015) for the respective compartments. **(d)** Depiction of the mode’s unique steady state solution (color coding as in (a)). Thick lines depict homeostatic cell numbers (i.e. the analytic steady state) for the estimated parameter set shown in (b). Thin lines represent time courses for arbitrary initial cell numbers. Initial values have been varied separately for each cell type, while leaving the other cell numbers at their respective steady state value.

Consequently, the subsequent downstream compartment, namely ST-HSCs, is described as follows:

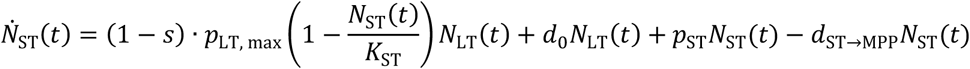

Herein, (1 – *s*) refers to the proportion of LT-HSCs which differentiate due to asymmetric cell division, while the influx of cells due to background differentiation of LT-HSCs is described by *d*_0_*N*_*LT*_(*t*). ST-HSCs proliferate with rate *p*_ST_ and differentiate with rate *d*_ST→MPP_.

Furthermore, in the MPP compartment, cells proliferate with rate an effective rate of 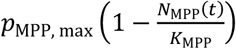 and differentiate with rate *d*_MPP→PB_ to further downstream compartments. *d*_ST→MPP_N_ST_(*t*) denotes the influx from the ST-HSC to the MPP compartment. The resulting ODE for the MPP compartment reads:

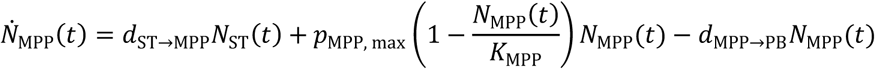

Further downstream compartments can be integrated by sequentially adding compartments as proposed in (Busch et al., 2015).

### Parameter estimation

In order to identify an optimal parameterization of the mathematical model, we apply a maximum likelihood method which minimizes the residual sum of squares (RSS) between the model simulation and the available data. In order to estimate cell turnover and differentiation, we use label progression data in a homeostatic system from (Busch et al., 2015). To appropriately account for this data type, we split each compartment in a labeled and an unlabeled fraction obeying the same parameters (see Supplementary Information for further details of the mathematical approach).

Since label progression data was acquired in young mice, we assume that about 80 % of LT-HSC divisions are asymmetric (Florian et al., 2018), thereby fixing the symmetry parameter *s* = 0.2 accordingly. Moreover, we adhere to estimates from (Busch et al., 2015) reporting that MPPs differentiate with rate *d*_MPP→CMP_ = 3.992 per day to the CMP compartment and with rate *d*_MPP→CLP_ = 0.022 per day to the CLP compartment, yielding a total MPP outflux of *d*_MPP→PB_ = 4.014 per day. As an upper limit for LT-HSC turnover, we fix *p*_LT, max_ = 1 per day. Due to the logistic growth limitation, this value is achieved only in the limiting case of an empty ST-HSC compartment and does not qualitatively influence our findings in steady state. We further make use of steady state compartment size ratios as estimated in (Busch et al., 2015), where 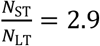 and 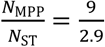. In our model formulation, these steady state compartment size ratios are independent from the carrying capacity *K*_ST_ (see Supplementary Information for further mathematical details). Therefore, we arbitrarily set *K*_ST_ = 10,000. Moreover, this allows us to reduce the set of parameters to be estimated to the LT-HSC background differentiation rate *d*_0_ and the effective proliferation rate of MPPs *p*_MPP, eff_ in steady state. Note that we only estimate a steady state value for the effective proliferation rate of MPPs here, since the label progression data was acquired in a homeostatic situation and therefore does not allow assessment of neither the maximum proliferation rate of MPPs *p*_MPP, max_ nor the respective carrying capacity *K*_MPP_. However, given a value for *p*_MPP, max_ and steady state values for *p*_MPP, eff_ and *N*_MPP_, a corresponding *K*_MPP_ can be calculated explicitly (see Supplementary Information for further details).

The table in Figure 1b summarizes both assigned and estimated parameter values.

## Results

### Formulation of a demand-driven LT-HSC model

We study control mechanisms for the most primitive HSC compartment (LT-HSC) within a mathematical modelling approach by assuming that (i) the turnover of and the outflux from the LT-HCS population is demand-driven and regulated at the progenitor level and (ii) LT-HSCs divide asymmetrically, while this ability declines with age. Technically, the model is implemented as a set of three, sequentially aligned compartments representing populations of LT-HSCs, ST-HSCs and MPPs (Fig. 1a, details in Methods). Assumption (i) is integrated as a logistic growth limitation of the LT-HSC compartment which is not regulated by the abundance of LT-HSCs but by the saturation of the downstream ST-HSCs. In homeostasis, this leads to a slow LT-HSC turnover as well as a low contribution to the subsequent compartments, while in perturbation scenarios, both LT-HSC turnover and differentiation are increased. The ability to undergo asymmetric cell division is incorporated as an explicit feature of LT-HSCs. Hence, the deregulation of asymmetric LT-HSC divisions with age (assumption (ii)) is modeled as an increase in the fraction of LT-HSCs that divide symmetrically rather than asymmetrically. ST-HSCs gain an influx from the upstream compartment and possess the ability to proliferate and differentiate, without any further regulation. MPPs also gain an influx from the upstream ST-HSCs compartment and differentiate. However, their proliferation is regulated by MPP demand in order to ensure a fast recovery after MPP depletion. The system can be tuned to achieve configurations in which all three cell populations are present and stably contribute over time.

### Applying the model to steady state hematopoiesis

In order to obtain a model configuration that applies to murine hematopoiesis, we fit our model to quantitative data on label progression in the homeostatic situation. In a reference experiment, Busch and colleagues *in vivo* marked a small number of LT-HSCs by using a YFP marker being almost uniquely inducible in LT-HSCs (Busch et al., 2015). Assessment of the temporal abundance of marked cells in the downstream compartments allowed to interrogate cell turnover and fluxes. Unlike the mathematical model in (Busch et al., 2015), which does not account for a regulated LT-HSC compartment, we here use this data to estimate parameters, such as proliferation and differentiation rates, of an advanced model that explicitly considers a regulation on the LT-HSC level. By using an optimization routine, we obtain values for all model parameters which are provided in the table in Fig. 1b. Fig. 1c illustrates that the parameterized model successfully describes the homeostatic situation in which the label slowly progresses from LT-HSCs to ST-HSCs and MPPs.

We estimate that LT-HSCs divide rarely, on average once per 111 days (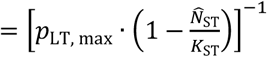). Similarly, background differentiation of LT-HSCs is a very rare event in homeostasis with an estimated value of once per 500 days (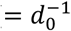). In contrast, both ST-HSCs and MPPs proliferate almost as fast as they differentiate.

### Modeling hematopoietic stress response

Unlike the homeostatic situation, hematopoietic recovery is mainly driven by LT-HSCs, which are activated in response to stress. Accordingly, depletion of downstream blood cells pushes LT-HSCs into cycle thereby increasing their differentiation and replenishing the system. Depletion of cells is commonly initiated by cytotoxic treatments or by irradiation followed by transplantation of donor cells whose contribution to hematopoiesis is subsequently measured (Essers et al., 2009; Osawa et al., 1996; Wichmann and Loeffler, 1985; Wilson et al., 2008; Wilson et al., 2009). However, the preconditioning does not only impact on hematopoietic cells but may also effect other tissues and thereby initiate further, indirect feedback loops that can influence hematopoietic recovery. Interestingly, Schoedel and colleagues established a perturbation model in which hematopoietic stem and progenitor cells were depleted *in situ* while minimizing the impact on other cell types (Schoedel et al., 2016). They could show that upon multilineage depletion, both ST-HSC and MPP populations quickly expanded close to steady state levels after few weeks only, whereas the LT-HSC compartment remained depleted for extended time periods.

In order to address the impact of different perturbation scenarios within our model system, we consider three scenarios: (i) multilineage depletion, (ii) progenitor (i.e. both ST-HSC and MPP) depletion, and (iii) LT-HSC depletion. We mimic those by adjusting the initial values for the respective compartments accordingly (Fig. 2).

**Figure 2.**
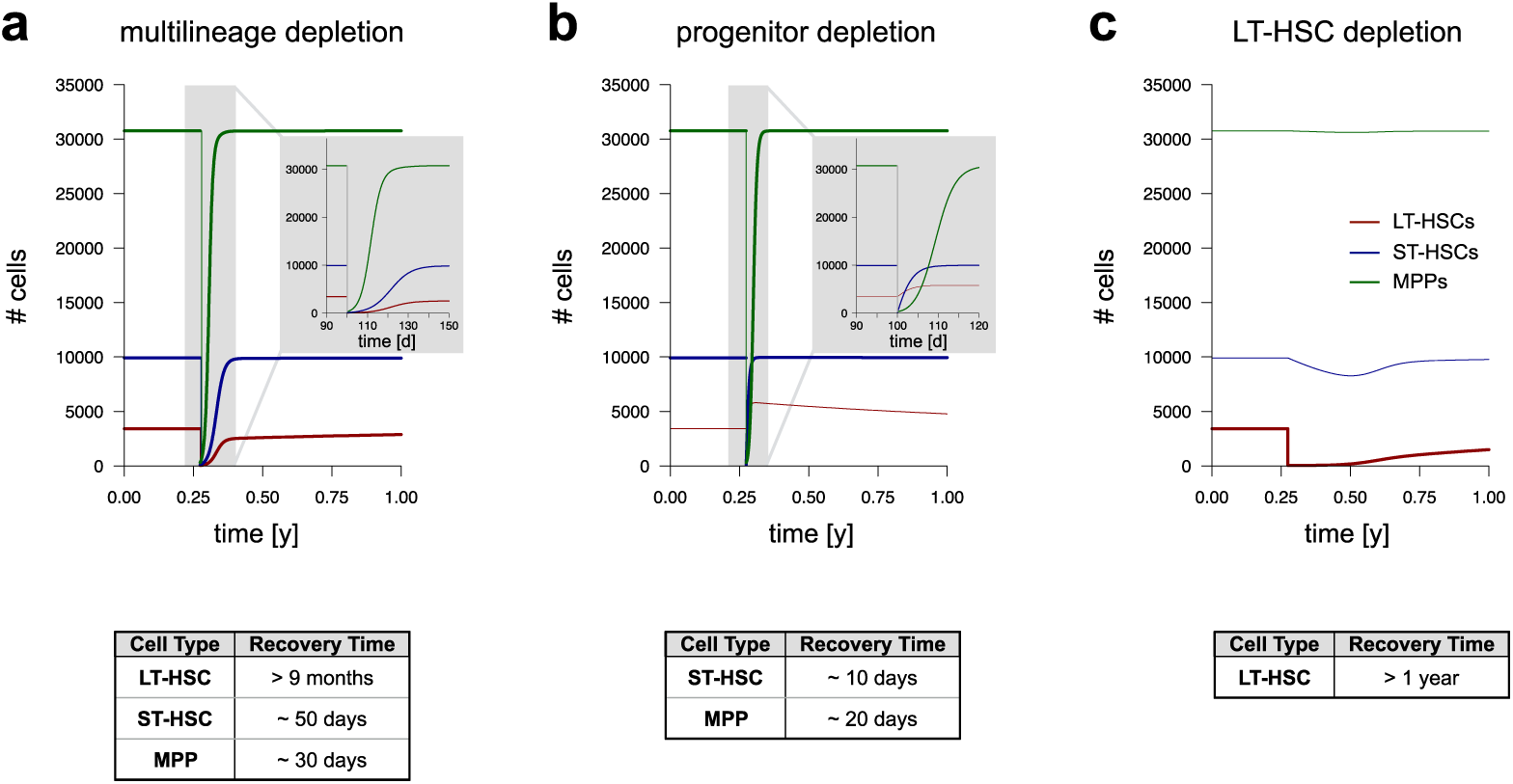
Hematopoietic stress response. Time courses illustrate the dynamic system response to targeted depletion (at *t* = 0.25*y*) of selected cell stages (depleting 99% of the cells): **(a)** Recovery of the system after almost complete depletion. Inset depicts a detailed view on the recovery of the system with time being given in units of days. **(b)** Recovery after both ST-HSC and MPP depletion. Inset allows a detailed view in time units of days. **(c)** Recovery after sole LT-HSC depletion. Time is given in units of years. The color scheme corresponds to the legend in (c). At time point zero, homeostatic cell numbers equal the steady state cell numbers obtained from Fig. 1d. Thin lines represent the time course of unperturbed compartments.

In the case of multilineage depletion (i.e. depletion of LT-HSCs, ST-HSCs and MPPs, which is comparable to the experiments conducted by Schoedel and colleagues) our model predicts a fast recovery of MPPs and ST-HSCs (Fig. 2a) to provide a sufficient supply of cells to peripheral blood. Here, the logistic growth limitation for the MPP compartment ensures a sufficiently fast MPP recovery after depletion: Due to an enhanced proliferation, MPPs go back to steady state levels within one month. Similarly, although being a little slower than MPPs, ST-HSCs reach steady state levels after 1 to 1.5 months. Interestingly, the model predicts that the regeneration of the LT-HSC pool is characterized by two phases: First, the LT-HSCs, which have remained after depletion, are forced into cycle due to ST-HSC demand resulting in a steep increase in cell numbers within the first weeks after depletion. When ST-HSCs have reached their steady state value, LT-HSC recovery slows down and does not reach the steady state level until more than 9 months.

Upon progenitor (both ST-HSC and MPP) depletion, both ST-HSC and MPP populations are predicted to recover after 10 to 20 days, as a consequence of LT-HSC proliferation being driven by ST-HSC demand and consequently increased differentiation (Fig. 2b). However, LT-HSC proliferation is also rapidly enhanced thereby satisfying the demand of the ST-HSC compartment. After this initial fast and strong increase, LT-HSC numbers only slowly return back to steady state levels, possibly due to low background differentiation.

In contrast to the fast recovery of hematopoietic stem and progenitor compartments after progenitor cell depletion, the model predicts a much slower return to the homeostatic situation, if only LT-HSCs are targeted. In fact, it is the almost stable and ongoing turnover and differentiation of ST-HSCs which does not pass on an increased demand to the LT-HSCs. Therefore, the population of LT-HSCs experiences a very slow recovery and only re-establishes over the course of more than 9 months (Fig. 2c).

In conclusion, our results indicate that the presented model successfully extends from the homeostatic situation to a qualitative and quantitative description of three reference scenarios associated with hematopoiesis under stress and regeneration. Our model extends these findings to the hypothesis that there are clear differences in the kinetics of LT-HSC and progenitor recovery after stress induction.

### Modeling hematopoietic aging

Several studies reveal that the abundance of phenotypic LT-HSCs increases with age, while the functional potential of these cells declines (Beerman et al., 2010; Dykstra et al., 2011; Florian et al., 2012; Geiger et al., 2013; Rossi et al., 2005). However, the reasons for the pronounced increase in LT-HSC number with aging are still not fully understood. It could be speculated that the low albeit constant demand of progenitor cells can only be met by a growing LT-HSC pool that compensates for their declining functionality. In this context, we recently discussed the limited ability of aged LT-HSCs to divide asymmetrically as a potential reason for their continuously increasing abundance (Florian et al., 2018). In fact, we observed about 80 % asymmetric divisions in LT-HSCs derived from young mice, while this number declines to about 20 % for aged donors. We raise the question whether this shift in the potential for asymmetric divisions is sufficient to explain the apparent increase in LT-HSCs.

For a quantitative assessment, we measured cell frequencies in LT-HSC, ST-HSC and MPP compartments in young (10-16 weeks old), middle-aged (40-54 weeks old) and old (more than 86 weeks old) C57BL/6 mice (Fig. 3a). LT-HSC frequencies increase almost exponentially, while ST-HSC and MPP frequencies remain rather constant or, in the case of MPPs, even tend to decrease over the mouse lifespan. For a comparison of the data with our model, we considered compartment size ratios which describe the compartment size relative to the abundance of LT-HSCs. We found that the ST-HSC to LT-HSC ratio rapidly decreases within the first year and afterwards remains constant, while the MPP to ST-HSC ratio slowly decreases with age (Fig. 3b). Since we assessed slightly different marker combinations to sort the HSC populations as compared to (Busch et al., 2015), we obtain different compartment size ratios (see Experimental Methods in Supplementary Information for further details).

**Figure 3.**
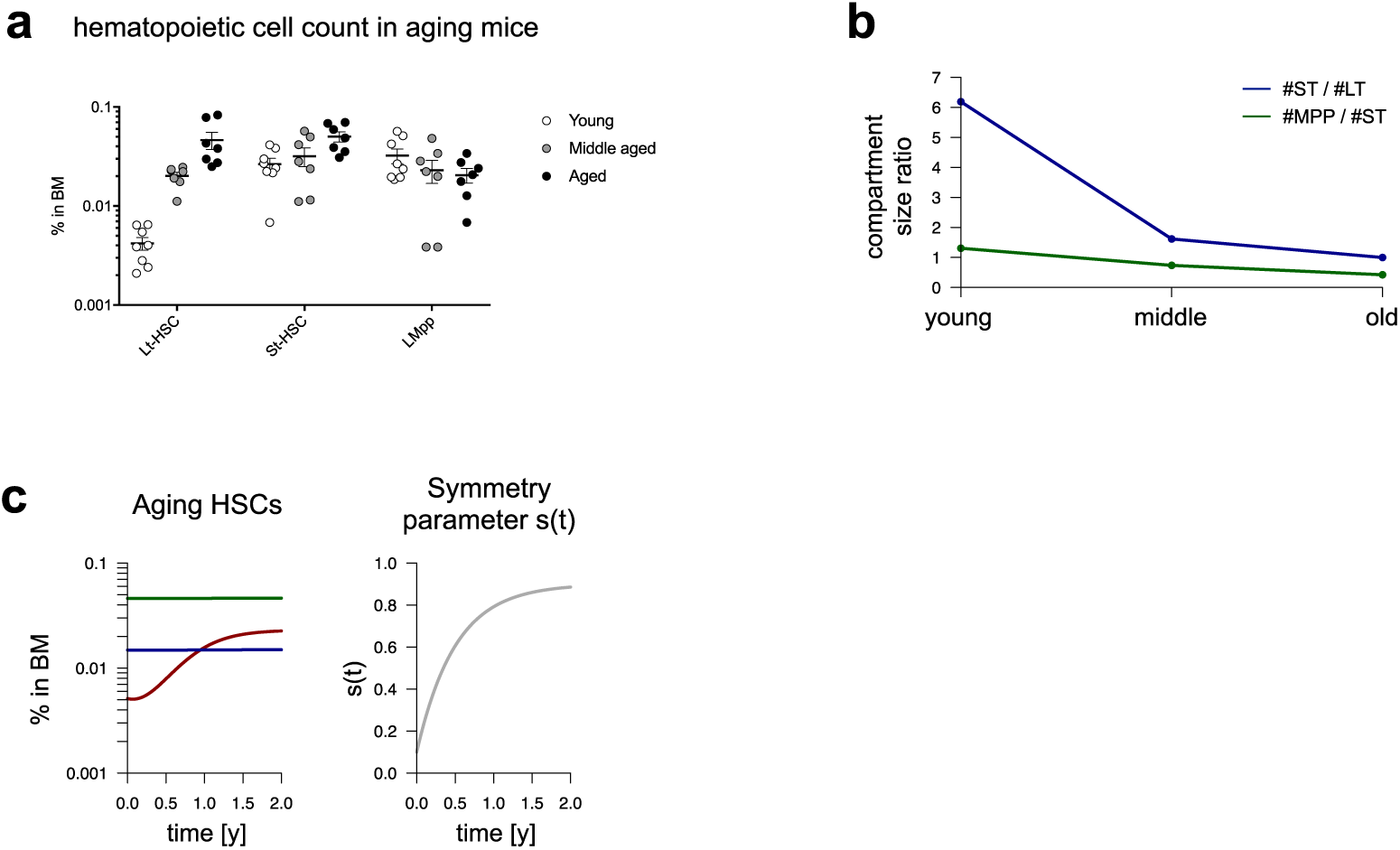
Hematopoietic aging. **(a)** Measured cell frequencies in LT-HSC, ST-HSC and MPP compartments for young (10-12 weeks), middle-aged (40-54 weeks) and old (more than 86 weeks) mice. Vertical lines represent the standard error of the mean of the measurements (see Supplementary Information for further details on cell sorting). **(b)** Measured compartment size ratios. **(c**) Cell frequencies predicted by the model (left-hand side) for a given change *s*(*t*) of the frequency of symmetric LT-HSC divisions with age (right-hand side, see Supplementary Information for further mathematical details). Cell frequencies are calculated by dividing the model solution by the mean total BM cell number which was used for cell sorting.

In our modeling approach, we implemented a corresponding aging scenario by assuming that the rate of symmetric cell divisions in LT-HSCs *s* = *s*(*t*) increases from 10% in a “young system” to 90% in an “old system”. Since the available data suggest a faster decrease in LT-HSC to ST-HSC ratio in the first compared to the second year, we chose a limited growth function for *s*(*t*), which increases rapidly within the first year and saturates within the second year (Fig. 3c, see Supplementary Information for mathematical details). By making these particular assumptions, we indeed see that the predicted frequency of LT-HSCs in BM strongly increases similar to the biological observations, whereas the frequencies of ST-HSCs and MPPs in BM remain rather constant (Fig. 3c). These findings illustrate that the increase in LT-HSC frequency may derive from a compensation mechanism, as with aging fewer LT-HSCs with asymmetric division contribute to the downstream ST-HSC compartment. The higher fraction of symmetric divisions expands the LT-HSC pool, but does not affect the outflux, which is only regulated by ST-HSC saturation level. Consequently, the model shows that there is no distinct change in the number of downstream cells. This confirms our experimental data and supports the prevailing notion that hematopoiesis in aged mice is still functional and does not suffer from severe insufficiencies in blood production.

To further use the capabilities of our modeling approach, we asked how a challenged hematopoietic system of an old mouse would recover after performing similar perturbation experiments as shown in the previous section, i.e. (i) multilineage depletion, (ii) progenitor (i.e. both ST-HSC and MPP) depletion, and (iii) LT-HSC depletion. Here, we defined an “old” mouse by assuming that the rate of symmetric LT-HSC divisions *s* would equal 0.8 while all other parameters remained unchanged. Thus, steady state LT-HSC numbers are increased compared to a young mouse.

In the case of multilineage depletion (i), the model predicts a very fast overshooting of LT-HSC numbers due to an interaction of enhanced proliferation driven by ST-HSC demand and predominant symmetric LT-HSC division (Fig. 4a). As soon as ST-HSC cell numbers get back to steady state levels, LT-HSC proliferation slows down, but LT-HSC numbers still need more than a year to get back to steady state levels – more time than the expected lifespan of a mouse at that age.

**Figure 4.**
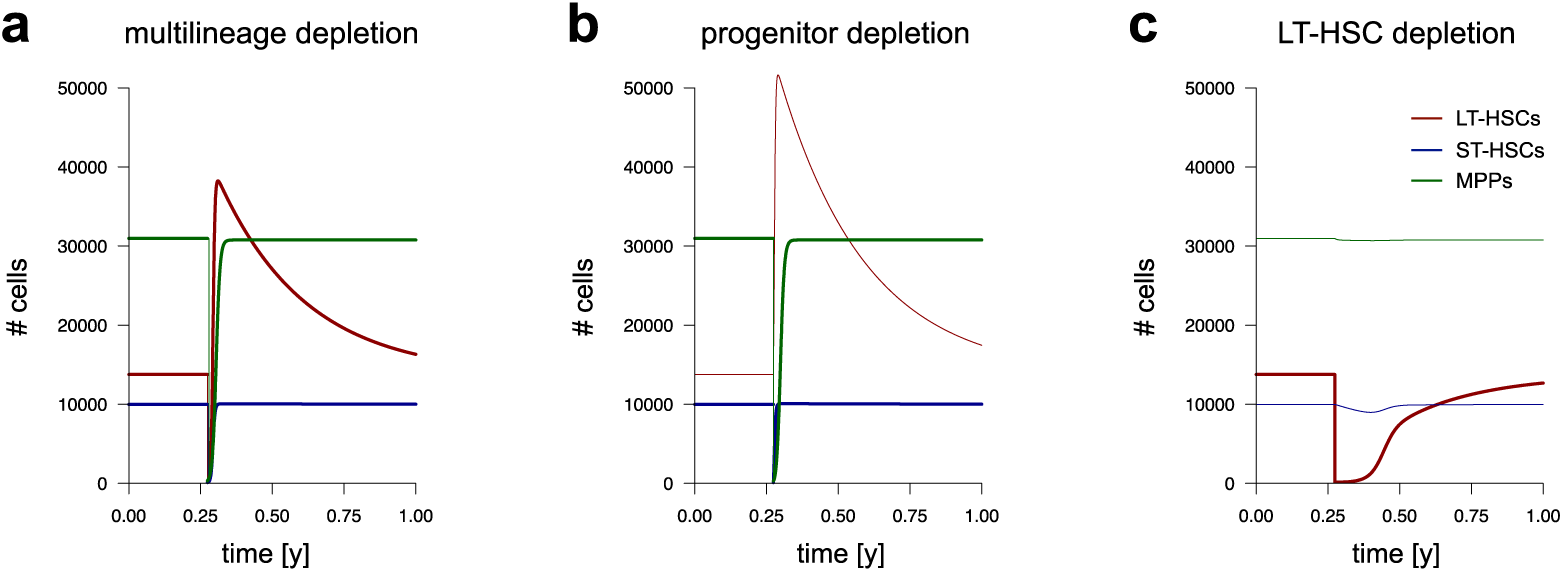
Hematopoietic stress response in an old mouse. Time courses illustrate the dynamic system response to targeted depletion (at *t* = 0.25*y*) of selected cell stages (depleting *99%* of the cells): **(a)** Recovery of the system after almost complete depletion. **(b)** Recovery after both ST-HSC and MPP depletion. **(c)** Recovery after sole LT-HSC depletion. Time is given in units of years. The color scheme corresponds to the legend in **(c)**. At time point zero, homeostatic cell numbers equal the steady state cell numbers for the symmetry parameter **s = 0.8**. Thin lines represent the time course of unperturbed compartments.

In contrast, both ST-HSC and MPP numbers are predicted show a similar behavior compared to the situation of a young mouse. Upon progenitor depletion (ii), the model predicts yet a higher overshooting of LT-HSC numbers, since in this scenario, LT-HSCs have not been depleted and progenitor recovery starts with LT-HSC numbers being in steady state (Fig. 4b). Again, progenitor recovery itself is comparable to the young situation, whereas LT-HSCs need more than a year to return to steady state levels.

LT-HSCs are predicted to recover faster (Fig. 4c) when compared to the young setting after sole LT-HSC depletion (iii). However, due to enhanced symmetric LT-HSC division and insufficient background differentiation, ST-HSC numbers decrease after some weeks thereby increasing LT-HSC proliferation. When

ST-HSC numbers saturate, LT-HSC numbers have already increased rapidly leading to a faster, albeit relatively slow, recovery of LT-HSCs compared to young mice.

Overall, the model predicts that an aged hematopoietic system will react very robustly to a major challenge, and presents with a general tendency of over-production of LT-HSCs, even in the phase of challenge.

## Discussion

Regulation of hematopoiesis and especially the role of LT-HSCs has been subject to intensive scientific debate for several decades (Busch et al., 2015; Laurenti and Gottgens, 2018; Moore and Lemischka, 2006; Morrison and Weissman, 1994; Rodriguez-Fraticelli et al., 2018; Schoedel et al., 2016; Sun et al., 2014; Weissman, 2000). While the definition of LT-HSCs stems from transplantation settings, recent findings on steady state hematopoiesis promote the concept that the LT-HSC pool is largely dispensable for supporting hematopoiesis in homeostatic conditions. With our mathematical modeling approach, we show that these different phenomena can be embedded in a quantitative concept in which LT-HSC response is driven by varying demands. The most prominent feature of our approach is to define HSC turnover not as a direct function of the LT-HSC population itself, but to attribute it to the on-demand regulation of a downstream compartment. Similar approaches have been pursued in other works, e.g. (Marciniak-Czochra et al., 2009), in which the most downstream compartment (or peripheral blood) is assumed to regulate both symmetry of division and proliferation of all upstream compartments. Although our assumption might oversimplify a more complex regulation, our approach emphasizes a regulatory control mechanism below the LT-HSC compartment that links continuous hematopoietic output with a rare activation of the LT-HSC reservoir. In fact, coupling LT-HSC turnover to the demand of ST-HSCs is sufficient to explain a diverse set of phenomena. Our results confirm the observation that LT-HSCs respond to a loss of more differentiated cell types with an increase in proliferation activity. Whenever downstream compartments have reached their respective steady state cell numbers, our modeling approach predicts decreasing proliferation of LT-HSCs irrespective of the number of LT-HSCs. Consequently, a sole depletion of LT-HSCs is not rapidly compensated, since no further demand is exerted from the downstream cell types. Hence, our model suggests that there is no or only a limited intrinsic regulation of the number of LT-HSCs solely based on their own abundance.

We further extended the model to account for changes in polarity and divisional asymmetry during hematopoietic aging. Based on our observation, that the polarity of Cdc42 and of correlated histone marks as well as the ability to divide asymmetrically are markedly reduced in LT-HSCs derived from older animals, we tested the hypothesis of an age-depended shift in the cells’ ability to divide asymmetrically as a critical feature explaining the LT-HSC pool expansion with aging. In contrast to the models presented in (Lander et al., 2009; Loeffler and Wichmann, 1980; Marciniak-Czochra et al., 2009), we explicitly neglected symmetric commitment of LT-HSCs, since we did not observe such divisions in our biological experiments. The results of our ODE model reproduced the aging-related increase of LT-HSC cell numbers, while ST-HSC and MPP numbers remained constant over time. Moreover, we were able to also quantitatively describe changes in the respective compartment size ratios. Taken together, our model supports the hypothesis that the aging-related shift from asymmetric to symmetric LT-HSC divisions is sufficient to describe phenomena of the aging LT-HSC compartment. Since there is a correlation between the loss of polarity (with respect to Cdc42 and correlated histone marks) in LT-HSCs and increasing numbers of symmetric divisions, our model emphasizes the idea that the shift from polar to apolar LT-HSCs is a major driver of hematopoietic stem cell aging.

In contrast to previous modeling approaches of the hematopoietic stem cell system by others (Abkowitz et al., 2000; Busch et al., 2015; Engel et al., 2004; Maloy et al., 2017; Manesso et al., 2013; Roeder et al., 2003) and us (Glauche et al., 2009), we here shift the focus from a self-sustaining and self-regulated LT-HSC population towards a population that is dynamically regulated by downstream compartments. Other than in the demand-driven model proposed in (Marciniak-Czochra et al., 2009), we place the focus on only LT-HSC proliferation being driven by ST-HSC demand rather than the most downstream compartment. Similarly, older model approaches discuss the role of asymmetric cell divisions in the context of balancing self-renewal and differentiation (Wichmann and Loeffler, 1985). In contrast to these works, we study the role of an impaired maintenance of divisional asymmetry during aging as functional/epigenetic mechanism to account for the increase in phenotypically defined LT-HSCs.

The generalizability of our model approach (as of any other) is on the one hand limited by the simplifications which are necessary to acquire identifiable model solutions, while on the other hand restrictions apply for the available data. As an example, one could argue that also different progenitor stages are sequentially involved in the underlying regulation of LT-HSCs or that there is an additional functional impairment of aged LT-HSCs. Moreover, we do not include any information on the contribution of the niche, which has been shown to also play a role in stress and homeostasis (Florian et al., 2018). However, without obtaining further data that investigates the system response to targeted, cell type specific perturbations, it is not possible to derive a more detailed understanding of further feedback regulations. As such, our focus is currently limited to the interplay between LT-HSCs and ST-HSCs, while the implemented MPP compartment is directly coupled to the ST-HSCs without further regulation.

Although our model approach adheres to the simplifying idea of reflecting different stages of HSC differentiation by the sequential coupling of intrinsically homogenous “compartments”, we understand this concept as a simplification of an inherently continuous process (see also (Laurenti and Gottgens, 2018) and references therein). In the experimental context, the phenotypic definition of cell types is a powerful tool to prospectively identify populations that are enriched for certain functional potentials and to allow setting up comparable experimental protocols. However, the objection that transits between the different cell stages are most likely not instantaneous but gradual also applies to the model context. Here, the formulation in terms of disjunctive, intrinsically homogenous cell compartments owes to the simpler mathematical formalisms and the easier comparison to experimental data.

Our approach illustrates that mathematical models are an essential tool to not only conceptually but also quantitatively embed a range of diverse biological phenomena in a unifying context. We demonstrate this potential by applying a novel model of HSC regulation to a range of observations in homeostatic situations that have challenged the prevailing view of an “almighty” stem cell. We conclude that a dynamically regulated and demand-driven approach is well-suited to explain hematopoiesis in the context of steady state blood production and response to stress and transplantation.

## Acknowledgment

We thank Thomas Höfer for providing published time course data from (Busch et al., 2015).

## Author Contribution

MK and IG conceived the mathematical model. MCF and HG designed and performed the experiments. MK and IG wrote the first draft of the manuscript and all authors approved the final manuscript.

## References

Abkowitz, J.L., Golinelli, D., Harrison, D.E., and Guttorp, P. (2000). In vivo kinetics of murine hemopoietic stem cells. Blood 96, 3399-3405.

Armanios, M., Alder, J.K., Parry, E.M., Karim, B., Strong, M.A., and Greider, C.W. (2009). Short telomeres are sufficient to cause the degenerative defects associated with aging. Am J Hum Genet 85, 823-832.

Beerman, I., Bhattacharya, D., Zandi, S., Sigvardsson, M., Weissman, I.L., Bryder, D., and Rossi, D.J. (2010). Functionally distinct hematopoietic stem cells modulate hematopoietic lineage potential during aging by a mechanism of clonal expansion. Proc Natl Acad Sci U S A 107, 5465-5470.

Busch, K., Klapproth, K., Barile, M., Flossdorf, M., Holland-Letz, T., Schlenner, S.M., Reth, M., Hofer, T., and Rodewald, H.R. (2015). Fundamental properties of unperturbed haematopoiesis from stem cells in vivo. Nature 518, 542-546.

Chambers, S.M., Shaw, C.A., Gatza, C., Fisk, C.J., Donehower, L.A., and Goodell, M.A. (2007). Aging hematopoietic stem cells decline in function and exhibit epigenetic dysregulation. PLoS Biol 5, e201.

de Haan, G., and Van Zant, G. (1999). Dynamic changes in mouse hematopoietic stem cell numbers during aging. Blood 93, 3294-3301.

Dykstra, B., Olthof, S., Schreuder, J., Ritsema, M., and de Haan, G. (2011). Clonal analysis reveals multiple functional defects of aged murine hematopoietic stem cells. J Exp Med 208, 2691-2703.

Engel, C., Scholz, M., and Loeffler, M. (2004). A computational model of human granulopoiesis to simulate the hematotoxic effects of multicycle polychemotherapy. Blood 104, 2323-2331.

Essers, M.A., Offner, S., Blanco-Bose, W.E., Waibler, Z., Kalinke, U., Duchosal, M.A., and Trumpp, A. (2009). IFNalpha activates dormant haematopoietic stem cells in vivo. Nature 458, 904-908.

Florian, M.C., Dorr, K., Niebel, A., Daria, D., Schrezenmeier, H., Rojewski, M., Filippi, M.D., Hasenberg, A., Gunzer, M., Scharffetter-Kochanek, K., et al. (2012). Cdc42 activity regulates hematopoietic stem cell aging and rejuvenation. Cell Stem Cell 10, 520-530.

Florian, M.C., Klose, M., Sacma, M., Jablanovic, J., Knudson, L., Nattamai, K.J., Marka, G., Vollmer, A., Soller, K., Sakk, V., et al. (2018). Aging alters the epigenetic asymmetry of HSC division. PLoS Biol 16, e2003389.

Geiger, H., de Haan, G., and Florian, M.C. (2013). The ageing haematopoietic stem cell compartment. Nat Rev Immunol 13, 376-389.

Geiger, H., Denkinger, M., and Schirmbeck, R. (2014). Hematopoietic stem cell aging. Curr Opin Immunol 29, 86-92.

Glauche, I., Moore, K., Thielecke, L., Horn, K., Loeffler, M., and Roeder, I. (2009). Stem cell proliferation and quiescence—two sides of the same coin. PLoS Comput Biol 5, e1000447.

Henry, C.J., Marusyk, A., and DeGregori, J. (2011). Aging-associated changes in hematopoiesis and leukemogenesis: what’s the connection? Aging (Albany NY) 3, 643-656.

Lander, A.D., Gokoffski, K.K., Wan, F.Y., Nie, Q., and Calof, A.L. (2009). Cell lineages and the logic of proliferative control. PLoS Biol 7, e15.

Laurenti, E., and Gottgens, B. (2018). From haematopoietic stem cells to complex differentiation landscapes. Nature 553, 418-426.

Loeffler, M., and Wichmann, H.E. (1980). A comprehensive mathematical model of stem cell proliferation which reproduces most of the published experimental results. Cell Tissue Kinet 13, 543-561.

Maloy, M., Maloy, F., Jakobsen, P., and Olav Brandsdal, B. (2017). Dynamic self-organisation of haematopoiesis and (a)symmetric cell division. J Theor Biol 414, 147-164.

Manesso, E., Teles, J., Bryder, D., and Peterson, C. (2013). Dynamical modelling of haematopoiesis: an integrated view over the system in homeostasis and under perturbation. J R Soc Interface 10, 20120817.

Marciniak-Czochra, A., Stiehl, T., Ho, A.D., Jager, W., and Wagner, W. (2009). Modeling of asymmetric cell division in hematopoietic stem cells—regulation of self-renewal is essential for efficient repopulation. Stem Cells Dev 18, 377-385.

Moore, K.A., and Lemischka, I.R. (2006). Stem cells and their niches. Science 311, 1880-1885.

Morrison, S.J., Wandycz, A.M., Akashi, K., Globerson, A., and Weissman, I.L. (1996). The aging of hematopoietic stem cells. Nat Med 2, 1011-1016.

Morrison, S.J., and Weissman, I.L. (1994). The long-term repopulating subset of hematopoietic stem cells is deterministic and isolatable by phenotype. Immunity 1, 661-673.

Noda, S., Ichikawa, H., and Miyoshi, H. (2009). Hematopoietic stem cell aging is associated with functional decline and delayed cell cycle progression. Biochem Biophys Res Commun 383, 210-215.

Osawa, M., Hanada, K., Hamada, H., and Nakauchi, H. (1996). Long-term lymphohematopoietic reconstitution by a single CD34-low/negative hematopoietic stem cell. Science 273, 242-245.

Rodriguez-Fraticelli, A.E., Wolock, S.L., Weinreb, C.S., Panero, R., Patel, S.H., Jankovic, M., Sun, J., Calogero, R.A., Klein, A.M., and Camargo, F.D. (2018). Clonal analysis of lineage fate in native haematopoiesis. Nature 553, 212-216.

Roeder, I., Loeffler, M., Quesenberry, P.J., Colvin, G.A., and Lambert, J.F. (2003). Quantitative tissue stem cell modeling. Blood 102, 1143-1144; author reply 1144-1145.

Rossi, D.J., Bryder, D., Seita, J., Nussenzweig, A., Hoeijmakers, J., and Weissman, I.L. (2007). Deficiencies in DNA damage repair limit the function of haematopoietic stem cells with age. Nature 447, 725-729.

Rossi, D.J., Bryder, D., Zahn, J.M., Ahlenius, H., Sonu, R., Wagers, A.J., and Weissman, I.L. (2005). Cell intrinsic alterations underlie hematopoietic stem cell aging. Proc Natl Acad Sci U S A 102, 9194-9199.

Schoedel, K.B., Morcos, M.N.F., Zerjatke, T., Roeder, I., Grinenko, T., Voehringer, D., Gothert, J.R., Waskow, C., Roers, A., and Gerbaulet, A. (2016). The bulk of the hematopoietic stem cell population is dispensable for murine steady-state and stress hematopoiesis. Blood 128, 2285-2296.

Sheikh, B.N., Yang, Y., Schreuder, J., Nilsson, S.K., Bilardi, R., Carotta, S., McRae, H.M., Metcalf, D., Voss, A.K., and Thomas, T. (2016). MOZ (KAT6A) is essential for the maintenance of classically defined adult hematopoietic stem cells. Blood 128, 2307-2318.

Sudo, K., Ema, H., Morita, Y., and Nakauchi, H. (2000). Age-associated characteristics of murine hematopoietic stem cells. J Exp Med 192, 1273-1280.

Sun, J., Ramos, A., Chapman, B., Johnnidis, J.B., Le, L., Ho, Y.J., Klein, A., Hofmann, O., and Camargo, F.D. (2014). Clonal dynamics of native haematopoiesis. Nature 514, 322-327.

Weissman, I.L. (2000). Stem cells: units of development, units of regeneration, and units in evolution. Cell 100, 157-168.

Wichmann, H.E., and Loeffler, M. (1985). Mathematical Modeling of Cell Proliferation: Stem Cell Regulation in Hemopoiesis, Vol 1 (Boca Raton, Florida: CRC Press, Inc.).

Wilson, A., Laurenti, E., Oser, G., van der Wath, R.C., Blanco-Bose, W., Jaworski, M., Offner, S., Dunant, C.F., Eshkind, L., Bockamp, E., et al. (2008). Hematopoietic stem cells reversibly switch from dormancy to self-renewal during homeostasis and repair. Cell 135, 1118-1129.

Wilson, A., Laurenti, E., and Trumpp, A. (2009). Balancing dormant and self-renewing hematopoietic stem cells. Curr Opin Genet Dev 19, 461-468.

Yu, V.W., Yusuf, R.Z., Oki, T., Wu, J., Saez, B., Wang, X., Cook, C., Baryawno, N., Ziller, M.J., Lee, E., et al. (2016). Epigenetic Memory Underlies Cell-Autonomous Heterogeneous Behavior of Hematopoietic Stem Cells. Cell 167, 1310-1322 e1317.

